# Matrix Gla Protein deficiency reveals a previously unrecognized mucosal calcification in the mouse airways

**DOI:** 10.64898/2026.11.07.765032

**Authors:** Elodie Baptista, Juliana Marulanda Montoya, Chaohua Deng, Lina Tabcheh, Romain Hugon, Joseph Deering, Tabea Kraft, Jean-Yves Jouzeau, Arnaud Bianchi, Marc D. McKee, Monzur Murshed, Hervé Kempf

## Abstract

Tracheal calcification is a common finding that is considered to have limited clinical significance when detected most often in elderly individuals. Despite its frequent occurrence, the cellular and molecular mechanisms at the origin of tracheal calcification remain largely unexplored. In this context, we aimed to characterize the development of tracheal calcification, and to determine the role of matrix Gla protein (MGP), a major inhibitor of ectopic calcification whose genetic or pharmacologic deficiency is associated with premature tracheal calcification in humans.

We found that wild-type mice undergo an unexpected early physiologic calcification of the tracheal cartilage rings beginning around one month after birth. This calcification progresses in a dynamic rostro-caudal sequence through a process involving terminal differentiation of tracheal chondrocytes. These findings challenge the prevailing view that tracheal cartilage is a permanent hyaline cartilage and instead reveals an early and highly organized physiologic calcification process. Furthermore, our findings further identify MGP as a key regulator of calcification within the respiratory tract. Indeed, MGP deficiency not only markedly accelerates tracheal cartilage calcification through a similar pattern and mechanism but also induces a previously unrecognized calcification of the tracheal mucosa, a lesion absent in wild-type mice.

These alterations could partly explain the respiratory manifestations observed in MGP-deficient patients having Keutel syndrome, which include dyspnea, cough, stridor, and recurrent infections, symptoms previously being mainly attributed to calcification of the tracheal cartilage rings.

## Introduction

Tracheobronchial calcification is a well-recognized radiological finding, yet it remains frequently overlooked in clinical practice and thus poorly studied and understood. Under physiologic conditions, calcification of the trachea and proximal bronchi is predominantly associated with aging and is generally considered to be asymptomatic and of no or little clinical significance. Its prevalence and severity increase markedly with age, although estimates vary according to the population examined. Tracheobronchial calcification is rarely observed before the age of 40, and it appears more frequently in women. Imaging studies have reported calcification in nearly 20% of elderly individuals^1^, including a radiologic series in which 19% of 105 elderly subjects displayed subtle-to-severe calcifications of the tracheal and bronchial cartilage.^2^ Biochemical analyses further support the appearance of mineral deposits during aging, since there exist a progressive accumulation of phosphate and calcium within the tracheal wall over time,^3^ consistent with the formation of calcium-phosphate mineral deposits similar to that found in bone or cartilage or in other types of ectopic calcification. Although usually clinically silent, extensive calcification of the tracheobronchial tree in elderly individuals can occasionally be associated with respiratory manifestations such as stridor, wheezing and dyspnea,^1^ thus raising the question of its potential functional consequences when calcification becomes particularly severe.

Notably, tracheobronchial calcification can occur prematurely and more extensively in rare but very specific clinical settings. Extensive tracheobronchial calcification is a hallmark of patients undergoing long-term anticoagulation therapy with vitamin K antagonists, such as warfarin.^2,4–10^ Indeed, the percentage of tracheobronchial calcification is estimated to be 47% in adults receiving warfarin sodium compared to 19% in age-matched controls.^2^ Premature calcification can also occur in children and young individuals treated with warfarin, in whom otherwise it would be exceptional.^10^ Several case reports have described massive tracheobronchial calcification in children receiving long-term warfarin treatment after Fontan surgery for single-ventricle palliation^8^ or following mitral valve replacement.^4,6^ Similarly, patients with Keutel syndrome (KS), a rare autosomal recessive disorder with fewer than 50 cases reported,^11^ also display premature and prominent tracheobronchial calcification, sometimes associated with airway stenosis and respiratory distress symptoms including dyspnea, cough, stridor and even recurrent infections.^10,12–18^ Together, these observations strongly point to Matrix Gla Protein (MGP), a potent vitamin K-dependent inhibitor of ectopic calcification^11^, as a key regulator of tracheobronchial calcification, since its activity is impaired by Vitamin K antagonists^19^ and since loss-of-function mutations in the *Mgp* gene cause KS.^20–26^

Although tracheobronchial calcification has been documented in these aforementioned clinical settings, little is known about its natural course. In particular, when calcification normally begins, how it progresses along the respiratory tree, whether tracheobronchial chondrocytes undergo a specific differentiation program, and how MGP deficiency modifies this process have not been systematically investigated *in vivo*.

In the present study, we performed a comprehensive anatomical, multiscale structural and spatiotemporal analysis of tracheobronchial calcification in both aging wild-type (WT) mice and *Mgp*-deficient mice, a close phenocopy model of KS.^27–30^ Here we show that in wild-type mice, tracheal cartilage calcification is not a late degenerative event, but a very early and highly organized process associated with terminal chondrocyte differentiation which progresses in a rostro-caudal manner along the trachea before extending into the bronchi. This is occurring within a markedly compressed time frame as compared with what is observed in humans.^1–3^ Importantly, MGP deficiency further accelerates the onset of this otherwise conserved process, mirroring impaired MGP activity in humans on a much shorter timescale.^12–18,20–26^ Most strikingly, at the same time, MGP deficiency in mice also induces an additional and previously unforeseen site of calcification affecting extracellular matrix in the lamina propria of the tracheobronchial mucosa, a finding that may have important and yet unexplored implications for respiratory physiology.

Together, these findings redefine the natural history of tracheobronchial calcification in mice, together with expanding its biological scope beyond the cartilaginous rings by uncovering a previously unappreciated calcification process within the murine airways under the control of MGP.

## Material and methods

### Animals

C57BL6/J and BALB/cJ wild-type (WT) mice, originally supplied by Charles River (Charles River, France), were locally inbred. *Mgp^+/+^*, *Mgp^+/-^*, *Mgp^−/−^*, *Mgp^−/−^;SM22-Mgp*; *Mgp^−/−^;Col2-Mgp*, *Mgp^−/−^;Hyp* and *Mgp^−/−^; Eln^−/−^; hELN* mice were described elsewhere.^27,29–33^

C57BL6/J and *Mgp^+/+^*mice, both WT at the *Mgp* locus and sharing the same genetic background are herein distinguished solely based on their breeding origin with *Mgp^+/+^* mice derived from heterozygous intercrosses. All mice were housed in conventional animal rooms under constant humidity (55 ± 10%) and temperature (22 ± 2°C) in 12h light-dark cycle conditions. They were fed *ad libitum* with standard food diet (Scientific Animal Food & Engineering, France) and had free access to water. Mice were placed in reproductive conditions whenever necessary to maintain the colony or provide a sufficient number of mice for experiments. For all the following procedures, adult mice were euthanized with CO_2_ in an appropriate chamber (Minerva) and newborn or very young animals were euthanized by decapitation. All animal protocols were in accordance with the guidelines set by the relevant animal care committee.

### Whole-mount Alcian Blue (AB) and Alizarin Red (AR) staining

Euthanized mice were decapitated, skinned, and eviscerated from their internal organs except for their full-length respiratory tract (trachea, bronchi and lungs). The specimens were then fixed in ethanol 95% for at least 2 days until the day of the experiments. Collected specimens were incubated in Alcian Blue 8GX (AB, Sigma A3157) solution (0.03% AB, 80% of 96% EtOH, 20% acetic acid) for approximately 18 hours. AB solution was replaced with 95% EtOH solution for at least 6 hours followed by 2% KOH solution until tissues were cleared out. Specimens were transferred to Alizarin Red S (AR, Sigma A5533) solution (0.003% in KOH 1%) until dark staining of the skeletal bones. Staining was then stopped at the very same time for all specimens in 1% KOH-20% glycerol solution while agitating. To store specimens for long periods of time, they were passed through glycerol/ethanol solutions (50/50, 80/20) and photographed in 100% glycerol solution.

### Histology and in situ hybridization

Tracheae were collected promptly after animal euthanasia. They were fixed in sterile 4% paraformaldehyde solution (prepared in RNase-free PBS) overnight at 4° and kept in 70% EtOH for at least 24 hours at 4°C. Specimens were then dehydrated via routine procedures using a Tissue processor (Leica ASP 300S) and embedded in paraffin. Blocks were then sectioned with a manual rotary microtome (Leica RM 2135) and sections were positioned on Superfrost+ glass slides (Fisher, Germany). Images were taken with a Leica DMD 108.

For von Kossa/Alcian Blue staining, sections were dewaxed and rehydrated through standard procedure. They were incubated for 1 hour in 2% silver nitrate (Normapur, VWR) solution (prepared in deionized water) under the light of a 60-Watt lamp, washed twice in deionized water for 3 minutes, once in 1% acetone for 3 minutes, and finally incubated for 15 minutes in Alcian Blue solution (0.02% Alcian Blue, 70% ethanol, 30 ml acetic acid). They were then washed for 1 minute in 1% acetone, 2 minutes twice in water, dehydrated (3 minutes in 25% ethanol (EtOH), 3 in 50% EtOH and 3 in 75% EtOH), counterstained with 0.1 % eosine for 30 seconds, destained twice for 3 minutes in 100% EtOH, washed twice with xylene for 5 minutes and mounted using Petrex^®^.

For detection of elastin fibers, Resorcin–Fuchsin/Van Gieson staining was performed. Sections were dewaxed and progressively rehydrated through a series of ethanol solutions with decreasing concentrations, followed by rinsing in distilled water. To visualize elastic fibers, sections were immersed in resorcin–fuchsin staining solution in a Coplin jar. The Coplin jar was placed in a 500-mL beaker containing 200 mL of water and heated in a microwave for 45 s. Sections were then left immersed in the staining solution overnight at room temperature. The following day, excess resorcin–fuchsin stain was removed by a brief rinse with 95% ethanol. Sections were subsequently washed under running tap water for 5 min and rinsed in distilled water. Sections were counterstained with Van Gieson solution for 30 s, rapidly dehydrated through increasing concentrations of ethanol, cleared in toluene, and mounted using Pertex® mounting medium (Histolab).

Nonradioactive *in situ* hybridization (ISH) was performed on 7 µm-thick paraffin embedded longitudinal or transverse sections of mouse trachea specimens mounted on Superfrost+ glass slides (Fisher). Sections were subjected to acid hydrolysis (0.2 N HCl, 15 min), proteinase K treatment (5 μg/ml in PBS, 15 min), postfixation (4% PFA, 5 minutes) and acetylation (0.25% acetic acid, 15 min). Each of these steps was followed by two 5-minute washes with PBS. After the last PBS wash, slides were rinsed with dH_2_O and air-dried, before a 2-hour prehybridization step, at 65°C, in hybridization solution [50% formamide, 10 mM Tris (pH 7.6), 200 μg/ml Torula yeast RNA, 1xDenhardt’s solution, 10% dextran sulfate, 600mM NaCl, 0.25% SDS, 1 mM EDTA (pH 8.0)]. Hybridization with digoxygenin-labeled *Collagen II* (*Coll II*) and *Collagen X (Coll X*) RNA probes was performed overnight at 65°C. Post hybridization, slides were rinsed briefly in 5x SSC at 65°C, washed with 1x SSC, 50% formamide (65°C, 30 minutes), subjected to RNase A digestion to reduce nonspecific hybridization, and washed at increasing stringency with SSC buffers (final wash at 55°C with 0.2xSSC). Bound probes were detected with an alkaline phosphatase-conjugated anti-DIG antibody (Roche) and revealed with BM purple substrate (Roche).

### Micro-computed Tomography (µCT)

µCT scanning of mouse aortas was performed with a SkyScan model 1272 instrument (SkyScan) set at a resolution of 8.0μm and 0.25-mm Al filter. Image acquisition was performed at 40Kv and 250µA with a 0.4-degree rotation between frames. Adaptive thresholding was applied to trachea and aorta separately. µCT image processing and analysis was performed with version 2.2f of the manufacturer’s software (SkyScan).

### X-Ray Microscopy (XRM)

Tracheae from five-week-old C57BL6/J and *Mgp^−/−^* mice were immersed in 70% ethanol and transferred to small polypropylene tubes containing 70% ethanol for XRM. Two low-magnification scans were acquired for each trachea using a Zeiss Xradia Versa 520 operating at 65 kV and 5 W, using a 4X objective lens, proprietary LE2 filter, and 1.4 s exposure time across 2201 projections to obtain volumes with a voxel size of 4.15 μm. The two volumes for each trachea were imported into Dragonfly 3D World, registered using mutual information, and stitched to form a single 3D volume containing the trachea of interest. Calcified tracheal rings and cricoid cartilage were segmented using grayscale thresholding. The thickness of calcification in each tracheal ring was measured using the volume thickness mapping tool, and calcified cartilage volumes were measured using connected component analysis.

For each trachea, a higher magnification volume was obtained near the first and second tracheal rings using a voltage of 65 kV, power of 5 W, 20X objective lens, LE2 filter, 15 s exposure time, and 2251 projections to obtain a final voxel size of 0.78 μm. Tracheal mucosal calcification and tracheal cartilage calcification were segmented using a convolutional neural network in Dragonfly 3D World. A five-layered 2.5D UNet classifier was trained by input of three ground truth datasets – each containing nine 2D images segmented into a background class, a calcified mucosa class, and a calcified tracheal ring class. The neural network was trained using a patch size of 80, stride-to-input ratio of 0.60, mini-batch size of 64, and learning rate of 0.40 with Adadelta optimization. The classifier was then applied to the full imaging volume to visualize all calcifications of the mucosa and tracheal rings.

### Statistics

All experiments on mouse specimens were repeated at least 3 times from 3-10 mice of each genotype. All data are reported as means ± S.E.M. with statistical significance defined as p<0.05(*) using two-tailed distribution with equal variance student’s t-test evaluated with Prism6 software (GraphPad).

## Results

### Tracheal calcification in C57BL/6J wild-type mice follows a specific spatiotemporal pattern

To identify the precise spatiotemporal occurence of tracheal calcification in WT mice, we dissected the respiratory tract of C57BL/6J mice at different ages and sequentialy stained them with Alcian Blue (AB) and Alizarin Red (AR). As expected, an initial morphological analysis first revealed that newborn mice (P0 to P7) displayed no evidence of calcification in the airway cartilage, and the oldest (6-month-old) specimens examined exhibited strong calcification of the rings along the whole respiratory system (data not shown).

Thus, we undertook a more thorough investigation covering a narrower range of ages between P14 and P60. AR staining revealed that the thyroid cartilage of the larynx shows the first signs of calcification at P14 (data not shown). The calcification spreads to the entire laryngeal cartilage over time (Fig.1A), with the cricoid cartilage being one of the last laryngeal structures to mineralize at P30 (Fig.1A). With regard to the trachea itself, some spots of AR staining could be found in the very first rings of P28 mice, with extensive calcification of the trachea clearly initiating at P30 as the first 5 or 6 cartilage rings display strong AB staining (Fig.1A). Calcification slowly progresses to adjacent distal rings in a dynamic rostro-caudal direction to fully cover the whole trachea and even extend into the two main bronchi after 2 months (Fig.1A). Of note, there does not seem to exist any sexual dimorphism in the calcification process of the tracheal structures since we found a similar pattern of initiation and progression between individuals, regardless of their sex (Supplemental Fig.1A). Interestingly, the number of rings affected, and their degree of calcification observed at P30, were virtually uniform between littermates and moderately variable between distinct litters (data not shown). Similar results were also obtained in the BALB/cJ strain (Supplemental Fig.1B)

**Figure 1.**
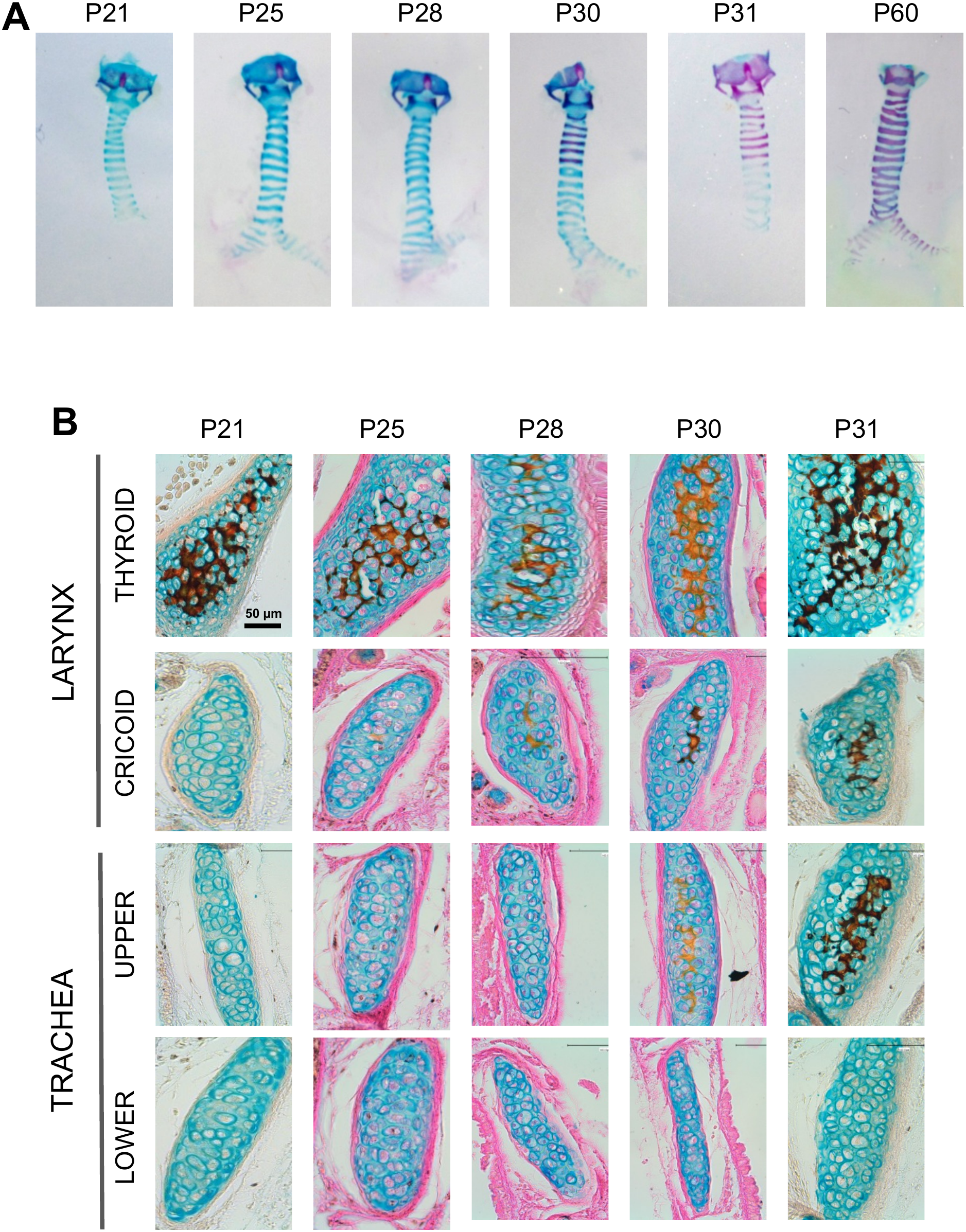
Kinetics of laryngeal and tracheal calcification in C57BL/6J mice. **(A)** Representative whole-mount Alcian blue/Alizarin red staining of the larynx and trachea of C57BL/6J mice at P21, P25, P28, P30, P31, and P60. Calcification is first detected in the thyroid cartilage and progressively extends throughout the larynx and along the trachea. At P30, calcification becomes visible in a few upper tracheal rings, whereas by P60, extensive calcification is observed throughout the trachea and main bronchi. **(B)** Representative longitudinal sections of the larynx and trachea stained with Alcian blue/von Kossa at P21, P25, P28, P30, and P31 (from left to right). Sections correspond to the thyroid cartilage (first row), cricoid cartilage (second row), upper (third row), and lower tracheal cartilage rings (fourth row). Calcified cartilage is identified by black/brown von Kossa staining. The thyroid cartilage is already markedly calcified at P21 and remains calcified at all subsequent time points. Calcification of the cricoid cartilage becomes detectable at P28 and increases thereafter. In the upper tracheal rings, calcification is first observed at P30 and becomes more pronounced at P31, whereas no calcification is detected in the lower tracheal rings over the same period.

Taken together, these morphological observations suggest that there is a very short period, around 4 weeks of age, when tracheal calcification initiates in C57BL6/J mice.

### Tracheal calcification affects the core of the cartilage rings

To validate and strengthen our morphological observations, histological experiments consisting of double staining with von Kossa/Alcian blue were performed on longitudinal (Fig.1B) and transverse (Supplemental Fig.2) sections of larynx and trachea from C57BL/6J mice at different ages.

**Figure 2.**
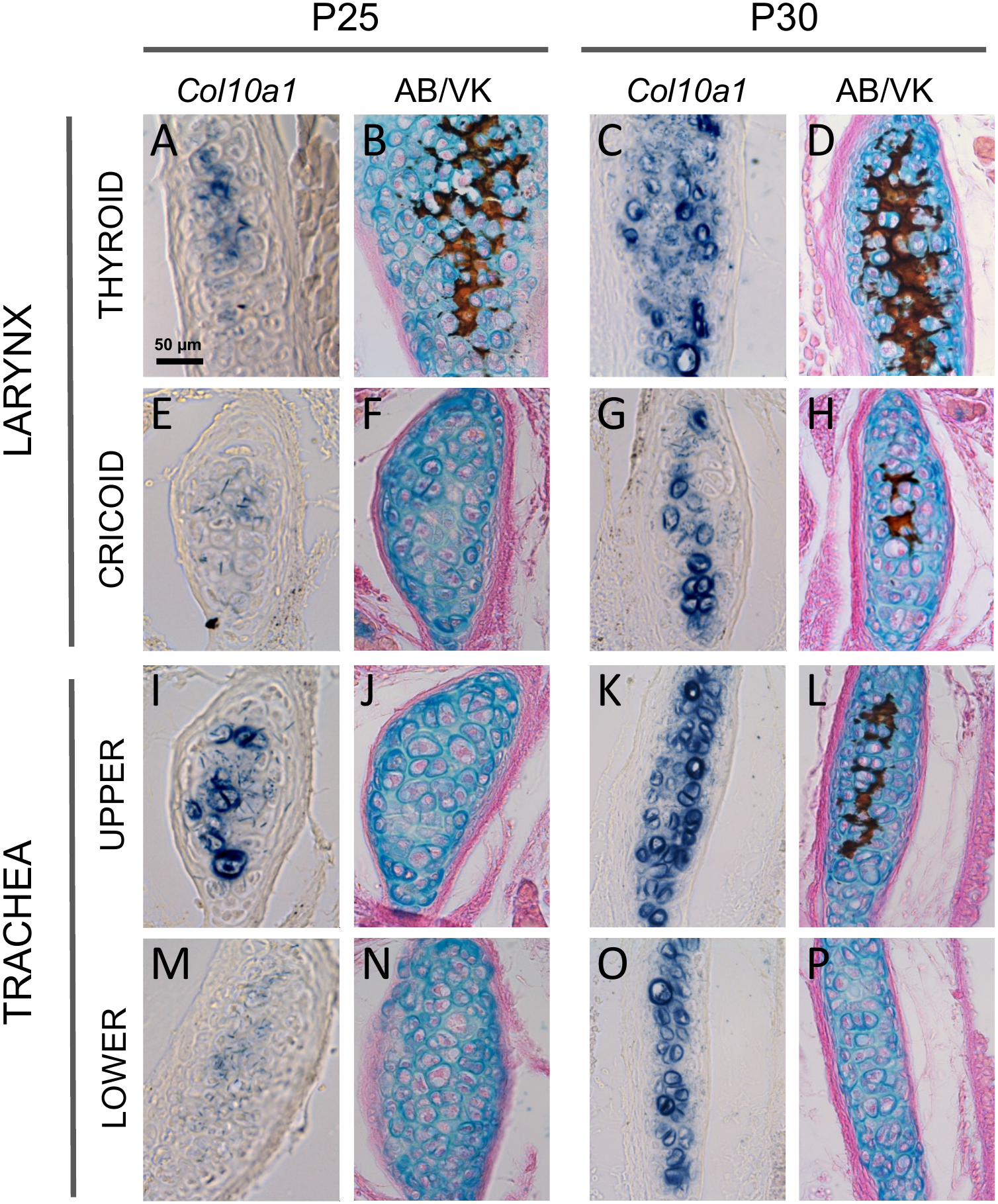
Spatiotemporal expression of Col10a1 along the laryngeal and tracheal cartilages. Longitudinal sections of the thyroid cartilage (**A-D**), the cricoid cartilage (**E-H**), a representative upper (**I-L**) and a lower tracheal cartilage ring (**M-P**) at P25 (**A-B, E-F, I-J and M-N**) and P30 (**C-D, G-H, K-L and O-P**). *Coll10a1* expression starts to be visible by *in situ* hybridization in all P25 samples (**A, E, I and M**) and is markedly induced at P30 in the larynx (**C, G**) and the trachea (**K, O**). Von Kossa staining appears as a time-dependent rostrocaudal pattern.

According to our morphological findings (Fig.1A), thyroid cartilage calcification can be noticed as early as P17 (data not shown) and spreads throughout the thyroid cartilage over time (Fig.1B and Supplemental Fig.2). Cricoid cartilage calcification can be observed starting at P28 (Fig.1B). Trachea calcification cannot be significantly detected by von Kossa staining in mice younger than 30-days old, neither in the upper part nor in the lower part (Fig.1B and Supplemental Fig.2). Starting at P30, calcification initiates in the upper part of the trachea and intensifies thereafter (Fig.1B). No calcification can be detected in the lower part of the trachea from P31 onwards (Fig.1B and supp Fig.2), although it fully calcifies later in time (data not shown and Fig.1A).

It is also important to note that longitudinal (Fig.1) and transverse (Supplemental Fig.2) histological sections also reveal that calcification initiates in the core of the cartilage rings, where chondrocytes become larger with time suggesting that they may undergo hypertrophy, and gradually spreads throughout the whole ring.

### Chondrocyte terminal differentiation precedes tracheal calcification

To verify that hypertrophy was taking place in cartilage rings over time, we looked at the expression of the specific marker of chondrocyte hypertrophy, *i.e. Col10a1* by *in situ* hybridization in the larynx and trachea of C57BL/6J mice at P25 and P30. *Col10a1*expression can be detected at low levels at P25 and at a strong level at P30 in longitudinal (Fig.2) and transverse (Supplemental Fig.3) sections of each region. Indeed, *Col10a1* can be detected at P25 in the thyroid region (Fig.2A) that is already calcified (Fig.2B), but it also can be found in regions that are not yet calcified, such as in the cricoid (Fig.2E-F) and tracheal rings (Fig.2I-J and Fig.2M-N), with a decreasing gradient between upper cartilage rings (Fig.2I) and lower cartilage rings (Fig.2M). At P30, *Col10a1* is strongly expressed in all respiratory tract regions analyzed (Fig.2C,G,K,O), even in lower cartilage rings that are still devoid of calcification at this stage (Fig.2P and supplemental Fig.3).

**Figure 3.**
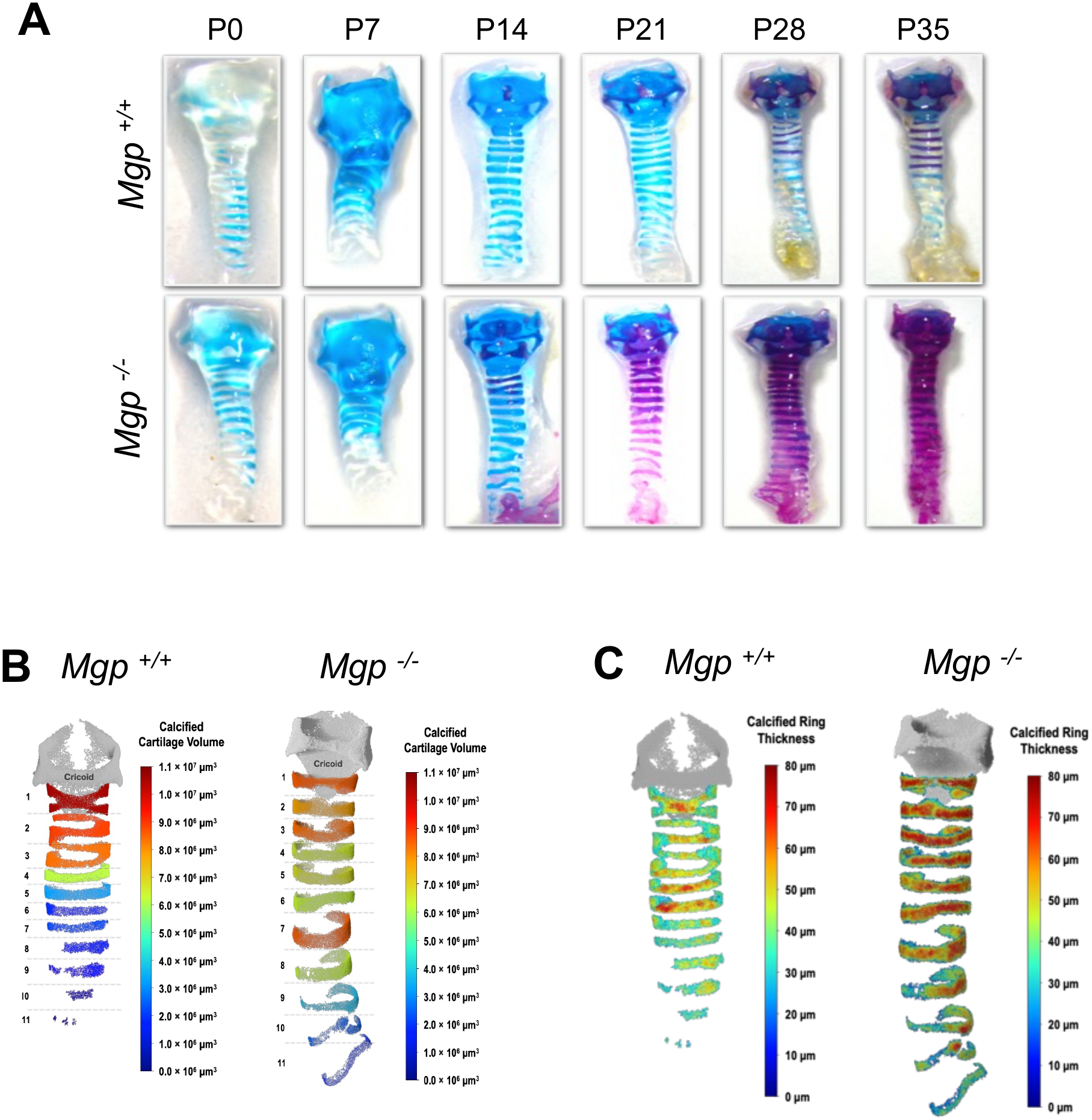
Accelerated and extensive tracheal calcification in Mgp-deficient mice. **(A)** Representative whole-mount Alcian blue/Alizarin red staining of the larynx and trachea from *Mgp*^+/+^ and *Mgp*^−/−^ littermates at P0, P7, P14, P21, P28, and P35. In *Mgp*^+/+^ mice, mineralization progressively develops from the laryngeal cartilages toward the trachea. In contrast, calcification occurs earlier and progresses more rapidly in *Mgp*^−/−^ mice, with calcification already visible in the upper tracheal cartilage rings at P14 and extending throughout the trachea by P21. **(B)** X-Ray Microscopy analysis and 3D reconstruction of tracheae from 5-week-old *Mgp*^+/+^ and *Mgp*^−/−^ mice showing the calcified cartilage volume of individual tracheal rings. Calcified cartilage volume follows a rostrocaudal gradient and is increased in *Mgp*^−/−^ compared with *Mgp*^+/+^ trachea. **(C)** Three-dimensional mapping of calcified tracheal ring thickness in 5-week-old *Mgp*^+/+^ and *Mgp*^−/−^ mice. *Mgp*^−/−^ mice display increased calcified ring thickness compared with *Mgp*^+/+^ mice.

### MGP-deficiency accelerates calcification of the tracheal cartilaginous structures

To investigate the potential role of MGP in mouse tracheal calcification, we next compared its spatiotemporal progression between *Mgp^+/+^*, *Mgp^+/−^* and *Mgp^−/−^* mice (Fig.3). Littermate *Mgp^+/+^* control mice on a pure C57BL/6J background display a pattern of tracheal calcification analogous to that of C57BL/6J WT mice (compare Fig.3A with Fig.1A). There was also no obvious difference in the tracheal calcification pattern in *Mgp^+/−^* mice heterozygous for the mutation (data not shown).

In contrast, MGP*-*deficient mice showed an earlier calcification of the trachea. Indeed, in these mice, calcification initiated in the upper tracheal rings at P14, 2 weeks earlier than in the *Mgp^+/+^* mice (Fig.3A). The spread of cartilage ring calcification followed a rostro-caudal progression identical to that observed in control mice (C57BL6/J or *Mgp^+/+^* mice) which completed in week’s time, at P21 (Fig.3A), when it typically requires about a month in *Mgp^+/+^*(Fig.3A)*, Mgp^+/−^* (data not shown) and C57BL/6J mice (Fig.1).

These morphological results were confirmed by high-resolution X-ray microscopy (submicron X-ray microcomputed tomography) analysis, which allowed quantification of the volume (Fig.3B) and thickness of calcified cartilage (Fig.3C) in the rings of 5-week-old *Mgp^+/+^* and *Mgp^−/−^* mice. These quantitative results demonstrate both an increasing rostro-caudal gradient across successive tracheal rings and, within each ring, a center-to-periphery gradient, confirming our AB-AR staining observations (Fig.3A). This approach also revealed a mild narrowing of the tracheal lumen in *Mgp^−/−^* mice compared to the more circular lumen observed in *Mgp^+/+^*mice (data not shown).

Comparison of *in situ* hybridization signals and von Kossa staining for mineral confirmed that, like in C57BL6/J mice (Fig.2), hypertrophy of tracheal chondrocytes precedes calcification, with upper rings being calcified before lower rings (Fig.4A). At P21, a stage where calcification is absent from the entire respiratory tract of *Mgp^+/+^* mice yet covers the full respiratory tract of *Mgp^−/−^* mice, *Col10a1* mRNA expression is higher when calcification is already engaged (Fig.4A). Of note, *Col10a1* expression is confined to the core of the rings that initiate hypertrophy and is surrounded by nonhypertrophic *Col2a1*-positive chondrocytes (Fig.4A).

**Figure 4.**
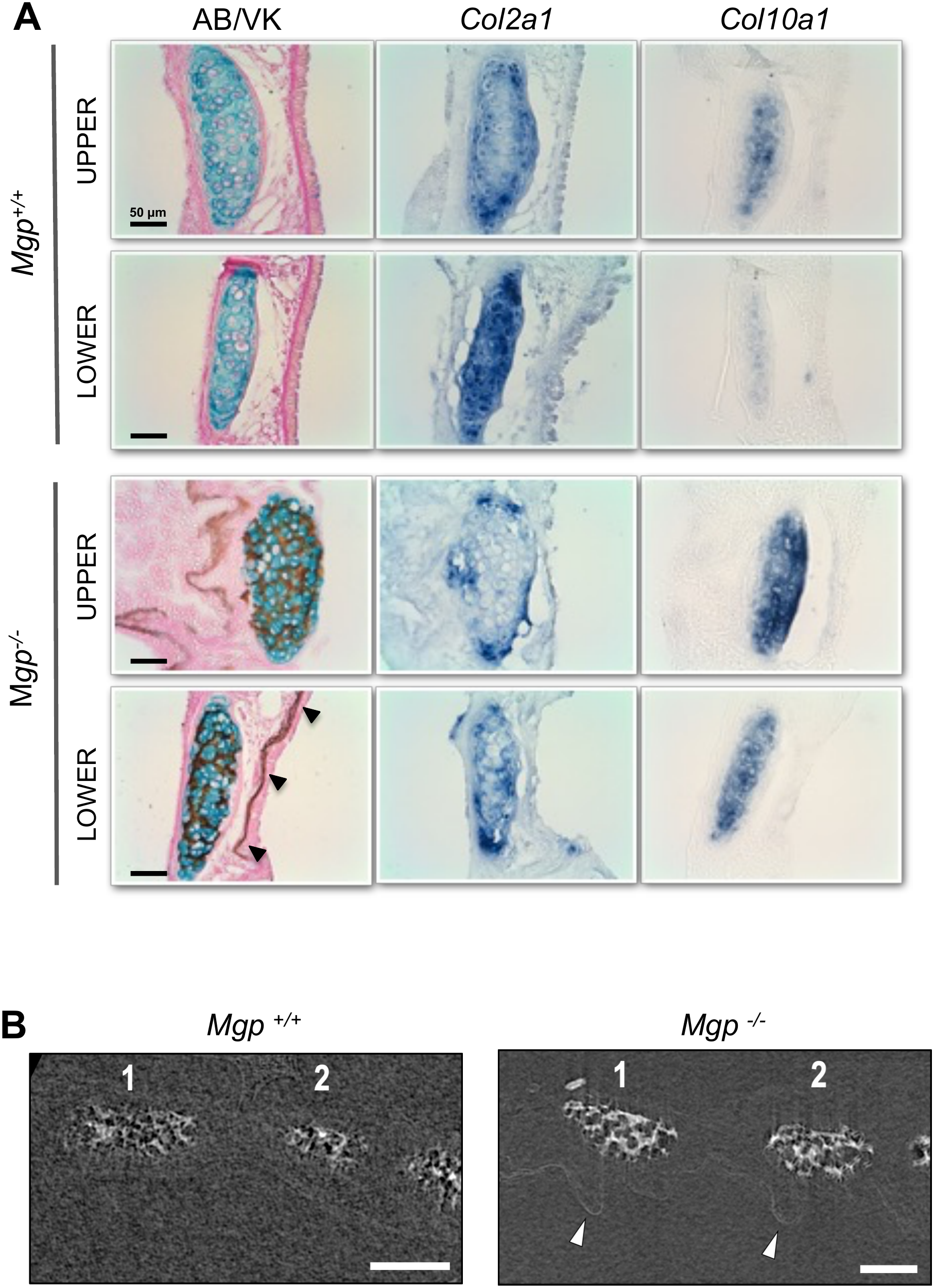
Absence of MGP accelerates cartilage ring calcification and induces calcification in the mucosa. **(A)** Representative Alcian blue/von Kossa staining and *Col2a1* and *Col10a1 in situ* hybridization of the trachea from *Mgp*^+/+^ and *Mgp*^−/−^ littermates at 3 weeks of age (P21). Increased cartilage calcification is observed in *Mgp*^−/−^ tracheae compared with *Mgp*^+/+^ controls. Black arrowheads indicate calcification extending beyond the cartilage into the surrounding mucosa. Scale bars are 50 µm. **(B)** Representative X-ray microscopy images of the upper trachea from *Mgp*^+/+^ and *Mgp*^−/−^ littermates. Tracheal cartilage rings 1 and 2 are indicated. White arrowheads highlight ectopic calcification within the tracheal mucosa of *Mgp*^−/−^ mice. Scale bars, 100 µm.

### MGP-deficiency induces extracartilaginous calcification in the trachea and the bronchi

Careful observation of *Mgp^−/−^* tracheal preparations revealed a diffuse additional calcification signal extending beyond the cartilaginous rings on the lumenal side of the trachea (Fig.3A) from P21 onward. This extracartilaginous staining was entirely absent at all stages in *Mgp^+/+^* (Fig.3A and white arrowheads in Fig.5A) and was also not observed in C57BL/6J mice, including at P60 when physiologic tracheobronchial cartilage calcification is maximal and extends throughout the respiratory tract (Fig.1A). This indicates that the diffuse calcification is specifically associated with MGP deficiency rather than with another acceleration of physiologic cartilage calcification.

**Figure 5.**
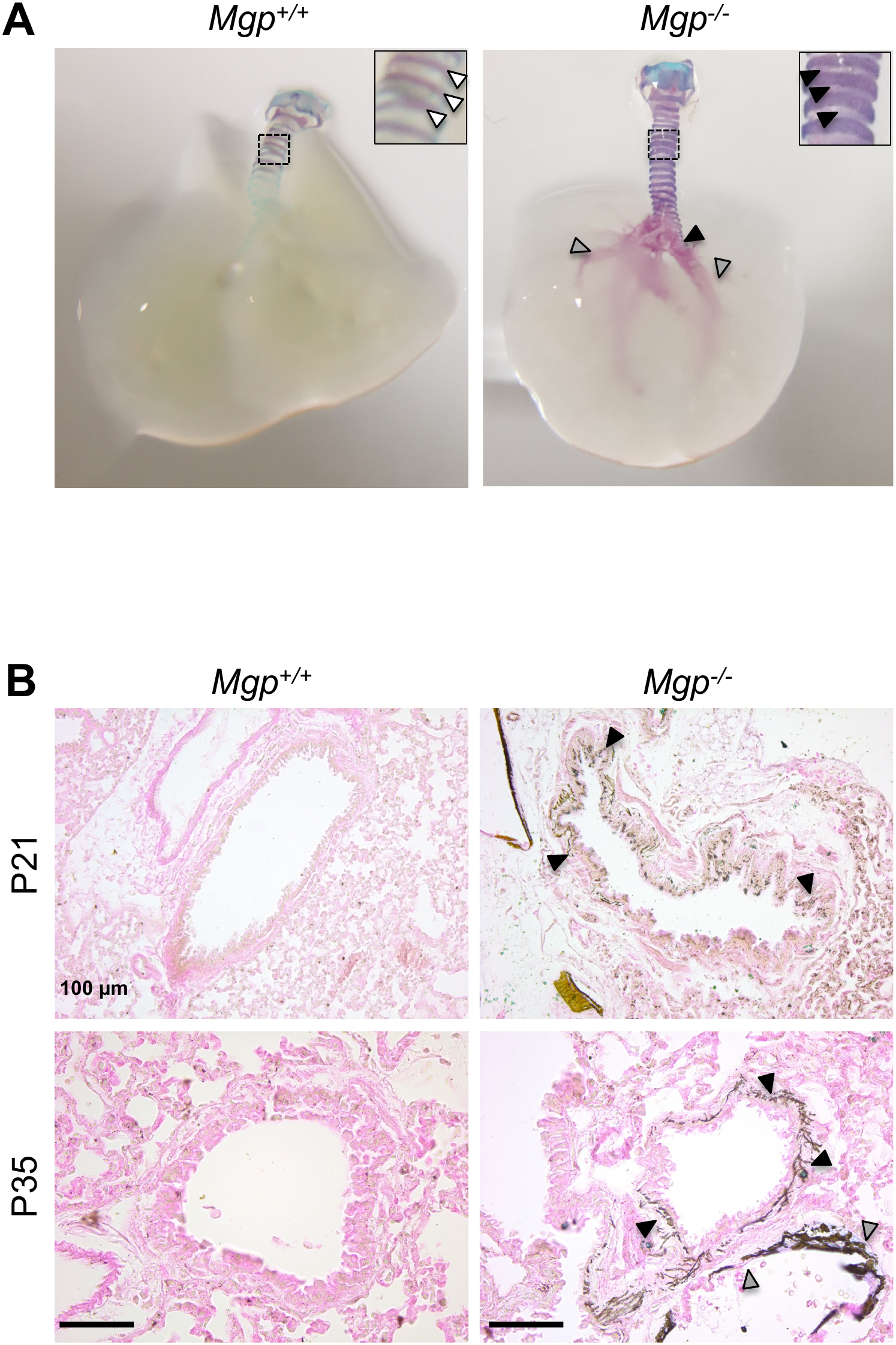
Progressive bronchial calcification in Mgp-deficient mice. **(A)** Representative whole-mount Alcian blue/Alizarin red staining of the larynx, trachea, and bronchi from *Mgp*^+/+^ and *Mgp*^−/−^ littermates at P35. White arrowheads show an absence of AR staining in between cartilage rings, where black arrowheads show positive staining. Grey arrowheads show calcification of lung arteries. **(B)** Representative Alcian blue/von Kossa staining of bronchial sections from *Mgp*^+/+^ and *Mgp*^−/−^ littermates at P21 and P35. Whereas no bronchial calcification is detected in *Mgp*^+/+^ mice, brown/black von Kossa-positive deposits reveal calcification of the bronchial wall in *Mgp*^−/−^ littermates (black arrowheads), in which there is also calcification in the media of pulmonary arteries (grey arrowheads). Scale bars, 100 µm.

Von Kossa staining for mineral confirmed that, in addition to the pronounced calcification of tracheal cartilage rings, *Mgp^−/−^* mice developed extensive mineral deposition within the tracheal mucosa (Fig.4A at P21 and Fig.6A at P35). High-resolution X-ray microscopy further confirmed the presence of this previously unrecognized tracheal mucosal calcification. Whereas C57BL6/J tracheae lacked any detectable extracartilaginous calcification, tracheae from 5-week-old *Mgp^−/−^*mice exhibited small but clearly detectable calcified deposits within the mucosal compartment (arrowheads in Fig.4B and supplemental video). Together, these complementary approaches demonstrate that MGP deficiency induces a distinct form of tracheal calcification affecting the mucosa in addition to cartilage.

**Figure 6.**
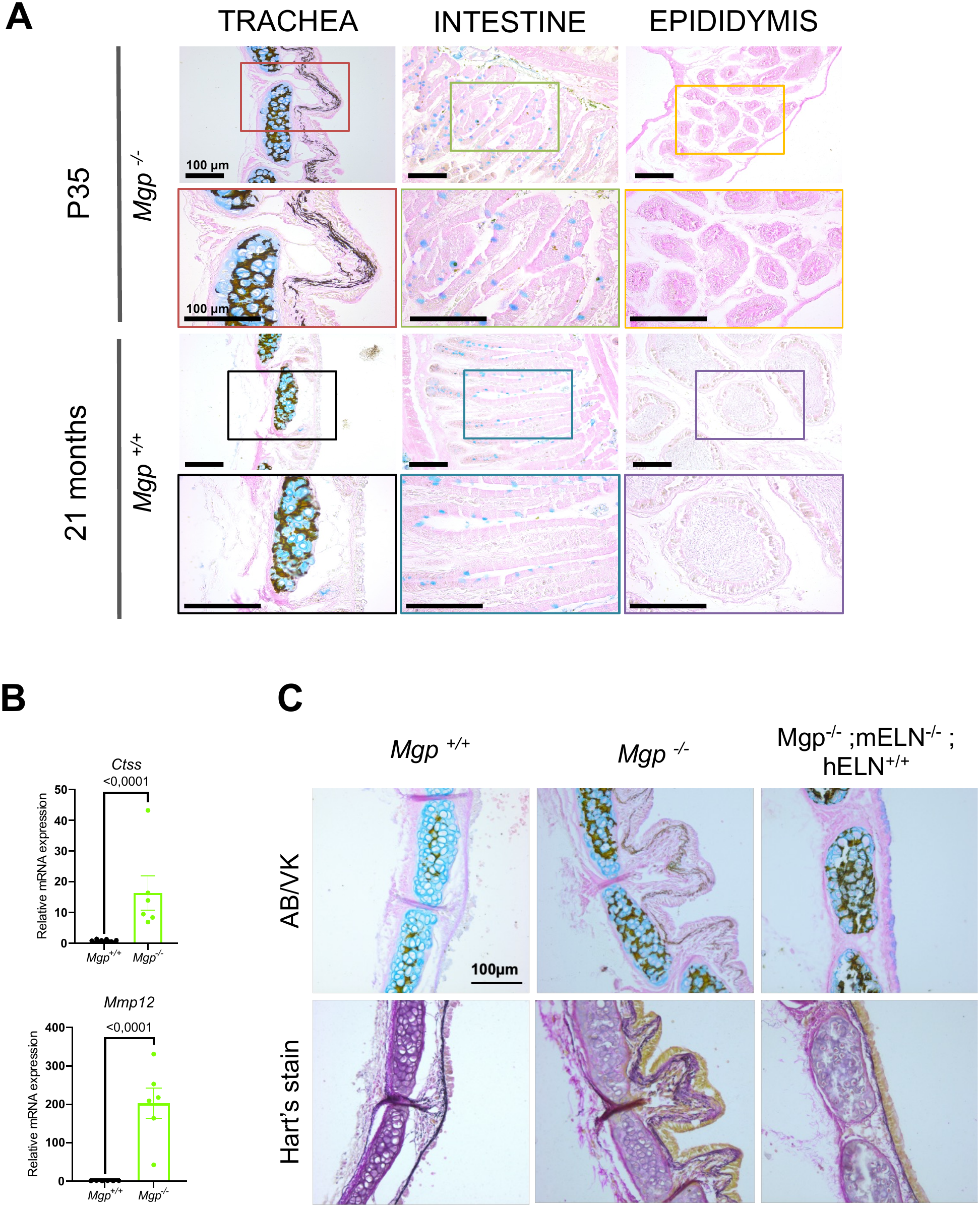
Unexpected mucosal calcification in the trachea of Mgp-deficient mice is associated with elastinocalcinosis. **(A)** Representative Alcian blue/von Kossa staining of the trachea, intestine, and epididymis from *Mgp*^−/−^ mice at P35 and *Mgp*^+/+^ mice at 21 months of age. Higher-magnification images of the boxed regions are shown below. Ectopic epithelial calcification is observed in the respiratory mucosa (lamina propria) of *Mgp*^−/−^ mice, whereas no mucosal calcification is detected in the intestine or epididymis. Scale bars, 100 µm. **(B)** Relative mRNA expression of *Ctss* and *Mmp12* assessed by RT-qPCR in *Mgp*^+/+^ and *Mgp*^−/−^ mice. Expression of both *Ctss* and *Mmp12* is significantly increased in *Mgp*^−/−^ compared with *Mgp*^+/+^ mice. Individual dots represent biological replicates (n=6); bars represent mean ± SEM. Statistical significance is indicated by the corresponding *P* values. **(C)** Representative Alcian blue/Von Kossa and Hart’s staining of tracheal sections from *Mgp^+^*^/+^, *Mgp*^−/−^ and *Mgp*^−/−^; *mEln*^−/−^; *hELN*^+/+^ mice. For the latter in which murine elastin deficiency is partly complemented by human elastin expression, elastocalcinosis is preserved and normal lamina propria structure is normal.

Importantly, this tracheal mucosal calcification was always associated with marked structural abnormalities of the epithelial lining. In *Mgp^−/−^* tracheae, the epithelium appeared tortuous, and markedly distorted, contrasting with the smooth and planar architecture of the noncalcified *Mgp^+/+^* mucosa (arrowheads in Fig. 4; Fig. 5B; Fig. 6A). High-resolution X-ray microscopy revealed that the mucosal calcified deposits followed a similarly wavy pattern, closely associated with the distorted epithelial architecture (arrowheads in Fig.4B and supplemental video). These observations suggest a close spatial relationship between mucosal mineral deposition and the profound structural alterations of the tracheal epithelium induced by MGP deficiency.

Moreover, mucosal calcification was not restricted to the trachea but extended into the bronchi, in which cartilage rings in mice is restricted to the most proximal branches.^34^ Except for the very few bronchial rings adjacent to the carina, no calcification was detected in *Mgp^+/+^* bronchi at P21 or P35 (Fig.5A-B) or at later stages (data not shown) through AR- or von Kossa stainings. In contrast, clear AR-positive staining was observed within the lungs of *Mgp^−/−^* mice, although the whole mount staining cannot totally distinguish between bronchial or vascular calcification (black and grey arrowheads respectively in Fig.5A-B). Histological analysis using von Kossa staining however clearly revealed calcification both within the bronchial mucosa (black arrowheads in Fig.5B) and pulmonary arteries known to be calcified in MGP-deficient mice (grey arrowheads in Fig.5B).^27,28^ Here again, like in the trachea, tortuous organization of the bronchial epithelium is occurring only when the mucosa is calcified in *Mgp^−/−^* mice (Fig.5B). The occurrence of mucosal calcification in noncartilaginous bronchial tissues further supports its distinction from cartilage-associated calcification and indicates that this pathologic calcification extends beyond the trachea along the conducting airways.

### Tracheobronchial mucosal calcification is MGP-deficiency specific and associated with elastocalcinosis

Because MGP deficiency markedly accelerates tracheobronchial cartilage calcification (hereafter noted as TCC), we next investigated whether tracheobronchial mucosal calcification (hereafter noted as TMC) might similarly represent an accelerated form of a physiologic aging process that could eventually occur in control animals. Strikingly, although TMC was fully established in P35 *Mgp^−/−^* mice (Fig.6A), it remained entirely absent in both *Mgp^+/+^* (Fig.6A) and *Mgp^+/−^* (data not shown) mice, even at advanced ages (21 months and later). Importantly, the marked epithelial distortion associated with TMC in *Mgp^−/−^* mice was likewise never observed in control animals, whose tracheobronchial epithelium retained a smooth and regular architecture (Fig.6A). Thess findings demonstrate that, unlike TCC, TMC and its associated epithelial structure alterations are not features of normal aging but rather a pathologic consequence of MGP deficiency.

To determine whether this phenotype reflected a broader susceptibility of epithelial tissues to calcification or was uniquely restricted to the respiratory tract, we examined additional organs with structurally distinct epithelial compartments, including the intestine and epididymis, known to express MGP.^35–37^ Neither intestinal nor epididymal tissues displayed detectable calcification or any sign of epithelial structural alteration both in *Mgp^+/+^* and *Mgp^−/−^* mice (Fig.6A), thus indicating that ectopic calcification associated with MGP deficiency does not broadly affect subepithelial compartments.

Given the high elastin content in the lamina propria of the tracheobronchial mucosa, histological analyses further suggested that TMC may mechanistically resemble the elastocalcinosis previously described in the vasculature of MGP-deficient mice.^27,38^ Indeed, von Kossa staining patterns indicated that mineral deposits closely associate with elastic fiber-rich regions and are accompanied by marked elastin fragmentation (Fig.6A), strongly suggesting that TMC develops through pathologic calcification of elastic structures. Consistent with this hypothesis, we observed by qPCR an upregulation of the mRNA expression of Cathepsin S (Ctss) and matrix metalloproteinase 12 (MMP12) (Fig.6B), two known elastolytic enzymes. These molecular signatures suggest that MGP-deficiency promotes a degenerative calcification process in which elastin remodeling and degradation create permissive conditions for mineral deposition.

To directly assess elastin’s contribution to TMC, we analyzed a novel compound mutant model (*Mgp^−/−^; Eln^−/−^; hELN*), recently reported,^33^ expressing approximately 60% less elastin than control mice. Remarkably, reducing elastin abundance drastically attenuated the progression and extent of TMC and the associated alteration of the epithelium structure (Fig.6C), demonstrating that elastic fibers are not passive but rather critical structural determinants required for TMC development.

### Differential rescue of tracheal cartilage and mucosal calcification in Mgp compound mutants

To further investigate the mechanisms by which MGP deficiency induces tracheobronchial calcification, we examined by µCT and histological staining TCC and TMC in three compound mouse models designed to restore MGP expression in distinct cellular compartments, or to reduce systemic phosphate (Pi) level (Fig.7).

**Figure 7.**
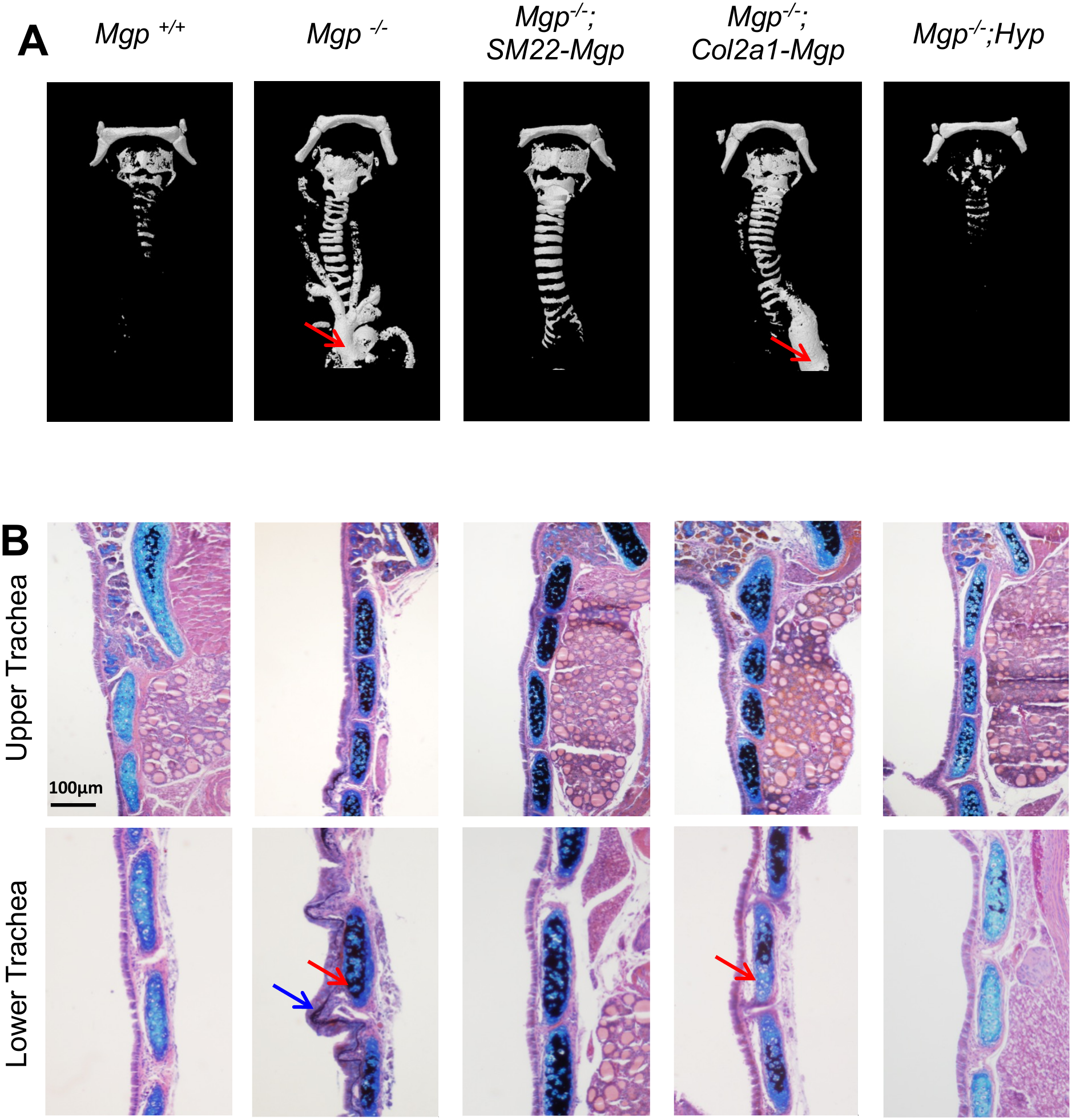
Micro-CT and histology comparing tracheal calcification in various compound mutants. **(A)** 3D micro-CT images of 3-week-old mice show comparable tracheal calcification in *Mgp^−/−^* and *Mgp^−/−^;SM22-Mgp* mice. There was an absence of vascular calcification in the later genotype compared to control (red arrow) but tracheal calcification is unaffected. In *Mgp^−/−^;Col2a1-Mgp* mice, calcifications were detected in both tracheal and vascular tissues. In contrast, *Mgp^−/−^;Hyp* mice show a reduction of tracheal calcification and a complete absence of vascular calcification. **(B)** Alcian blue/von Kossa staining of histology sections supports the µCT data. However, there was a clear indication (red arrows) that despite a weaker expression the *Col2a1-Mgp* transgene, there is some protective effect on tracheal calcification in *Mgp^−/−^;Col2a1-Mgp* mice. Interestingly, the calcified lamina propria (blue arrow) visible in *Mgp^−/−^* sections was not present in any of the compound mutants. Scale bar, 100 µm.

In *Mgp^−/−^*;*SM22-Mgp* mice, restoration of MGP expression driven by the smooth muscle-specific SM22 promoter completely rescues the extensive vascular calcification of *Mgp^−/−^* (Fig.7A), as previously shown.^31^ In contrast, this has no detectable effect on the accelerated calcification of the tracheal cartilage rings, which remains comparable to that observed in *Mgp^−/−^* mice (Fig.7A-B). Thus, although SM22-driven MGP expression is sufficient to prevent vascular calcification, it does not protect tracheal cartilage from calcification. We next examined *Mgp^−/−^*;*Col2-Mgp* mice, in which MGP expression is restored in chondrocytes and, as shown previously,^30^ has no protective effect on vascular calcification (Fig.7A). In contrast to the SM22-driven transgene, cartilage-specific MGP overexpression partially prevents the accelerated calcification of lower tracheal cartilage rings observed in *Mgp^−/−^*mice (Fig.7A-B). Interestingly, analysis of transgenic *Mgp* expression in different cartilaginous tissues revealed substantially lower expression in tracheal cartilage than in nasal cartilage for instance (Supplemental Fig. 4), which may, at least in part, explain the incomplete rescue of the tracheal phenotype. Finally, we examined *Mgp^−/−^;Hyp* mice. In these mice, in which systemic phosphate levels are lowered by approximately 50% through renal phosphate wasting,^32^ the TCC phenotype is reduced to a level similar to control mice (Fig. 7A), together with complete rescue of the vascular phenotype.^32^

With regard to TMC, it is first noteworthy that, in contrast to TCC, which was readily detected by conventional µCT, subepithelial calcification remained below the detection threshold of this approach (Fig.7A). This contrasts with the higher-resolution ability of submicron X-ray microscopy which was able to detect TMC (Fig.4B and supplemental video). Subsequent analysis of von Kossa-stained sections from the same animals allowed us to discern TMC and revealed that a different pattern emerges as TMC was absent in all three compound mouse models analyzed (Fig.7B). In *Mgp^−/−^*;*SM22-Mgp, Mgp^−/−^*;*Col2-Mgp and Mgp^−/−^;Hyp* mice, there was prevention of TMC accompanied by restoration of epithelial architecture with the tracheal lining presenting a smooth and regular organization resembling that of *Mgp^+/+^*mice (Fig.7B)

Taken together, these results show that TCC and TMC are differentially affected by local MGP expression and systemic phosphate reduction.

## Discussion

To investigate tracheal calcification *in vivo* in mice, we performed a study to define when and where calcification occurs, and we examined the mechanisms that regulate this process during normal aging and in the context of MGP deficiency.

### Tracheal cartilage calcification is an early physiologic event in mice

Our findings challenge the conventional view of tracheal cartilage as permanent, static hyaline cartilage by showing that TCC is an early physiologic event in mice. In C57BL/6J mice, TCC begins around 4 weeks after birth in the proximal tracheal rings and subsequently follows a dynamic rostro-caudal progression, ultimately expanding within the entire trachea and proximal bronchi by 2 months. A similar spatial pattern was observed in BALB/cJ mice, although with a slightly earlier onset, indicating that the timing of calcification varies slightly between strains while its overall progression is similar.

Given the extensive use of mouse models in research, it is surprising that early physiologic TCC has, to our knowledge, not been reported in mice. Our results, however, agree and thus extend and strengthen, older but sparse observations in birds and mammals. In the early 1980’s, Hoggs characterized mineralization of the syringeal, laryngeal and tracheal cartilages in birds and reported for that the first tracheal ring starts to mineralize 126 days post-hatching.^39^ Interestingly, besides the shape (complete in birds) and the number of rings (>100), the avian TCC differs from that we observed in mice in several respects: *i*) it follows a caudo-rostral pattern, *ii*) it remains incomplete at the cranial end, and *iii*) it undergoes ossification.^39^ In mammals, TCC has been described in otters and bats, with ring calcification detected in bats as early as 37 days and developing in a cranial-to-rostral progression.^40^ In rats, three reports by Bonucci *et al*. in 1974^41^ and Sasano *et al.* in the mid-1990’s^42,43^ likewise indicate that tracheal cartilage calcifies in relatively young individuals. These studies documented von Kossa staining at 10 weeks and appearance of hypertrophic chondrocytes in the central region of the rings as early as 4 weeks. This sequence is generally consistent with our observations in mice, although we detected von Kossa- and Alizarin Red-positive staining earlier in the upper tracheal cartilage of P30 C57BL6/J mice. So, among rodents, there seems to be a clear shift towards early stages for the physiologic calcification of tracheal cartilage in mice versus rats. However in both rodents, the process involves chondrocyte hypertrophy and cartilage calcification,^42,43^ rather than the ossification described in birds.^39^ At the molecular level, herein, we observed a progressive upregulation of *Col10a1* mRNA in the upper and lower tracheal rings, further supporting terminal chondrocyte differentiation. This localization differs partly from the findings of a previous rat study,^43^ in which type X collagen immunoreactivity was confined to the uncalcified peripheral region of the trachea rather than the mineralized or hypertrophic zones at 4,8 and 10 weeks.^43^

While the physiologic role of TCC in rodents remains unclear, several roles have been proposed in other vertebrate species. In birds, it has been associated with vocalization.^39^ In bats it relates to the development of flight and echolocation and is thought to protect the airways against deformation during rapid changes in thoracic and subglottic pressure.^39^ In diving otters, TCC has similarly been proposed to prevent airway collapse under increased external pressure.^40^ Whether physiologic TCC provides somehow comparable mechanical adaptation in mice remains to be determined.

#### MGP restrains the timing of tracheal cartilage calcification

MGP has long been recognized as a potent inhibitor of ectopic calcification in vascular and cartilage tissues.^11^ In addition to extensive arterial calcification, *Mgp*-knockout mice were originally reported to develop abnormal premature tracheal calcification based on observations made in a 5-week-old animal.^27^ Because our study shows that physiologic TCC begins around 4 weeks of age, we challenged this conclusion and revisited this interpretation by systematically comparing the tracheal phenotypes of *Mgp^+/+^*, *Mgp^+/−^* and *Mgp^−/−^* C57BL/6 mice through detailed multiscale analyses. Our data show that calcification is not normally detected during the first 4 weeks of life in *Mgp^+/+^* mice but is already evident 2 weeks after birth in *Mgp^−/−^* mice. MGP deficiency therefore accelerates an otherwise physiologic TCC program rather than fundamentally changing its spatial pattern. Indeed, once initiated, calcification follows the same rostro-caudal progression seen in *Mgp^+/+^* and C57BL6/J mice, ultimately involving all tracheal rings and extending into the few bronchial rings. This accelerated phenotype is consistent with the premature airway calcification reported when MGP activity is impaired, including in Keutel syndrome^12–18,20–26^ and during warfarin exposure.^2,4–10^ These findings identify MGP primarily as a modulator of the timing of TCC. Notably, this effect appears far more pronounced in humans where impaired MGP activity can advance TCC by several decades, whereas MGP deficiency advances its onset by only about two weeks in mice.

### Tracheobronchial mucosal calcification is a distinct consequence of Mgp deficiency

A major and unexpected finding of the present study is the identification in *Mgp^−/−^* mice of a previously unrecognized form of calcification affecting the lamina propria of the tracheobronchial mucosa (TMC). This TMC was associated with remodeling of the mucosal compartment, including marked structural alterations of the overlying epithelium, which appeared highly tortuous, together with development of fibrosis and hyperplasia (data not shown). Not found in *Mgp^+/+^* mice, even up to almost 2 years of age, this indicates that, unlike TCC, TMC is therefore not part of the normal aging-related calcification of the murine respiratory tract but represents a distinct and direct pathologic consequence of complete MGP deficiency.

Another important aspect of our finding is that TMC could only be visualized in 3D using high-resolution sub-µCT imaging and remained undetectable by conventional µCT in mice. Thus, similar microscopic calcifications may have escaped detection in humans, because routine imaging identifies calcification of the tracheal cartilage but may lack the resolution threshold required to reveal mineral deposits within the mucosa. Whether TMC occurs in patients with Keutel syndrome or following inhibition of MGP activity during warfarin exposure, therefore remains an important unanswered question. This possibility is particularly relevant because patients with Keutel syndrome frequently present with dyspnea, cough, wheezing and recurrent respiratory infections, symptoms until now generally attributed to extensive TCC.^12–18^ Our findings in mice suggest that this interpretation may be incomplete. C57BL6/J mice develop extensive TCC early in life without obvious respiratory complications, and physiologic mineralization or even ossification of the tracheal cartilages also occurs early in bats or birds without pathologic implication but rather physiologic adaptations.^39,40^ These observations argue that TCC alone may therefore be insufficient to explain the respiratory manifestations associated with MGP deficiency as claimed in Keutel syndrome case reports.^12–18,20–26^

TMC may thus represent a previously overlooked contributor to respiratory dysfunction in Keutel syndrome patients. Ectopic calcification of the lamina propria, together with epithelial remodeling, could alter the functions of the overlying epithelium, including mucus homeostasis and mucociliary clearance, thereby contributing to cough and susceptibility to respiratory infections through reduction of airway defense. Although the short lifespan of *Mgp-/-* mice (around 2 months)^27^ limits long-term respiratory studies, the early onset of TMC makes this model possibly valuable for determining how TMC may affect epithelial organization and function.

### Tracheal compartments display distinct susceptibilities to calcification

The rescue experiments indicate that MGP deficiency affects the tracheal cartilage and mucosa through distinct regulatory mechanisms. TCC retains a strong local component, because chondrocyte-specific MGP expression (but not expression in SM22-positive cells) protects, although partially, the tracheal rings from accelerated calcification. The partial rescue may reflect the substantially lower transgene expression in tracheal cartilage than in nasal cartilage, where the same transgene completely prevents calcification.^30^ The marked reduction of TCC in *Mgp^−/−^;Hyp* mice further shows that this locally regulated process remains sensitive to systemic phosphate availability, as in the vascular compartment.^31^

TMC follows a different pattern. Its prevention by *Mgp* expression from either *Sm22*- or *Col2a1*-expressing tissue as well as by systemic phosphate reduction, suggest that protection of the mucosa does not require MGP production exclusively in lamina propria. MGP produced in adjacent tissues may contribute to the local inhibitory environment of the mucosa. The broad rescue of TMC and epithelial tortuosity across the three compound models is consistent with a predominantly non-cell-autonomous regulation.

An additional observation further supports the existence of compartment-specific susceptibility to calcification. Despite the extensive arterial smooth muscle calcification cause by MGP deficiency in mice,^27–29,31,33,38^ we did not observe calcification in the trachealis muscle in *Mgp^−/−^* mice. Because tracheal smooth muscle cells can calcify *in vitro* under high-phosphate conditions,^44^ this apparent resistance *in vivo* is unlikely due to an intrinsic inability to calcify. Although it remains to be investigated, it suggests that the loss of MGP is insufficient to induce alone calcification in every smooth muscle cell and that the local environment is an additional major determinant of calcification.

Altogether, our findings distinguish two forms of tracheal calcification. TCC is an early physiologic process in mice that shares important features with the calcification observed in humans, although it develops over a much shorter time frame. MGP deficiency accelerates this normal cartilage calcification program. In contrast, TMC is a distinct pathologic consequence of MGP loss that affects the tracheobronchial mucosa, is associated with epithelial distortion, and is regulated differently from TCC.

The identification of TMC may change how the respiratory complications of MGP deficiency are understood. If TMC is present in patients with Keutel syndrome, it could provide a new explanation for their respiratory manifestations and challenge the prevailing view that these symptoms arise solely from TCC. Because TMC may escape conventional imaging, similar mucosal calcification could have been overlooked during warfarin exposure or in other respiratory disorders. Further studies should therefore determine whether TMC occurs in these conditions and establish how it might affect epithelial organization, mucociliary clearance and/or airway defense. In other words, in addition to clarifying the role of MGP in TCC, our study identifies TMC as a previously unrecognized component of airway pathology with potential relevance to Keutel syndrome and beyond.

## Acknowledgments

We are very grateful to Lauralie Christophe and Sébastien Francois (IMoPA animal facility) for taking care of the mice. We thank the Integrated Quantitative Biology Initiative (IQBI, McGill University) for use of their equipment. We thank Comet Technologies Canada (Montreal) for the free-of-charge academic license for the use of Dragonfly 3D World software. M.D.M. is the Canada Research Chair in Biomineralization, and a member of the Fonds de Recherche du Québec-Santé (FRQS) Centre for Structural Biology Research at McGill University, and the National Network for Advancing Bone Health Research in Canada (OsNET). We also thank the members of all teams for their technical help and valuable discussions. M.M. and M.D.M. are both members of the FRQS Network for Intersectorial Research in Sustainable Oral and Bone Health. H.K is a member of the International Network on Ectopic Calcification (INTEC).

