## Supplementary figures and images for "Matrix Gla Protein deficiency reveals a previously unrecognized mucosal calcification in the mouse airways"

### Baptista et al Supp Figure1.pdf

**A**

Male Female

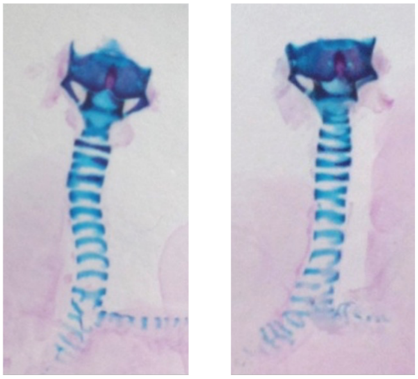

**B**

P21 P25 P27 P29 P30 P45

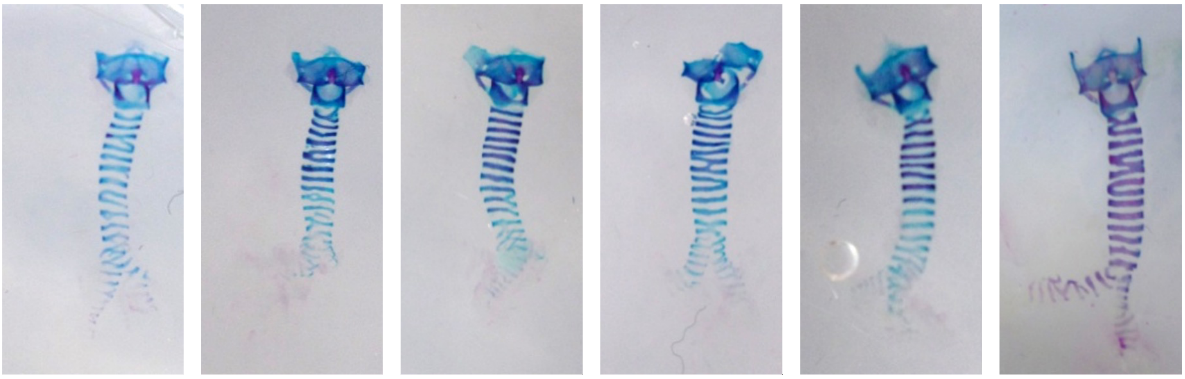

### Baptista et al Supp Figure2.pdf

**Supplemental Figure 2** **Baptista et al.**

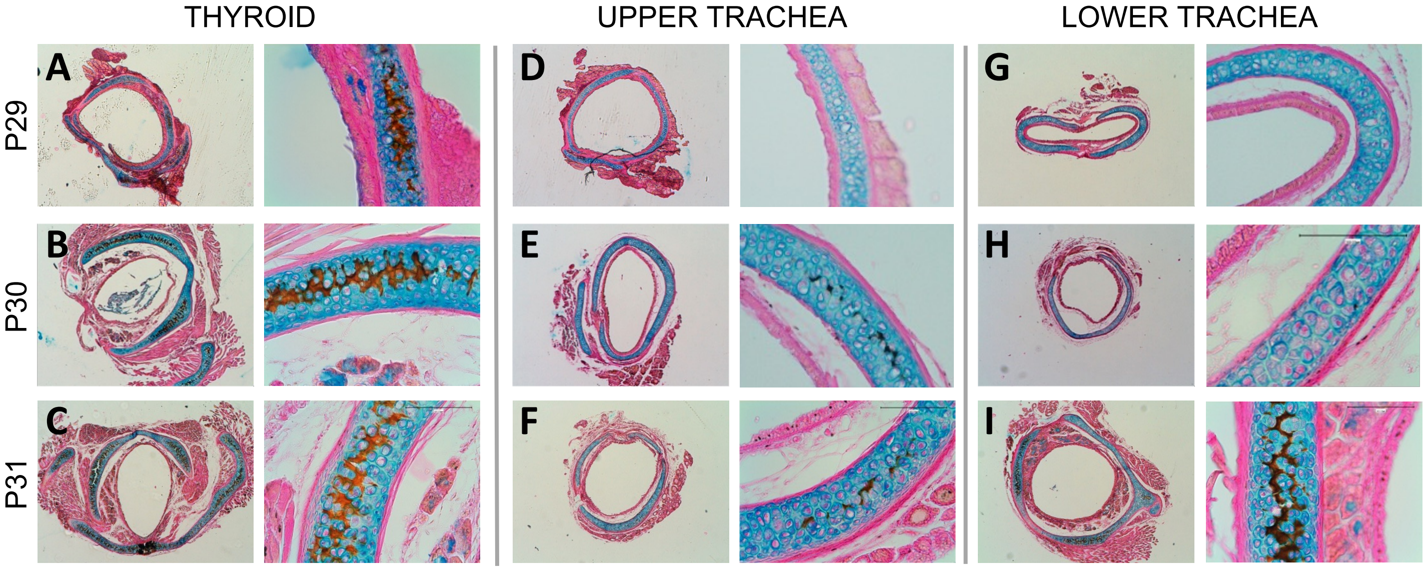

### Baptista et al Supp Figure3.pdf

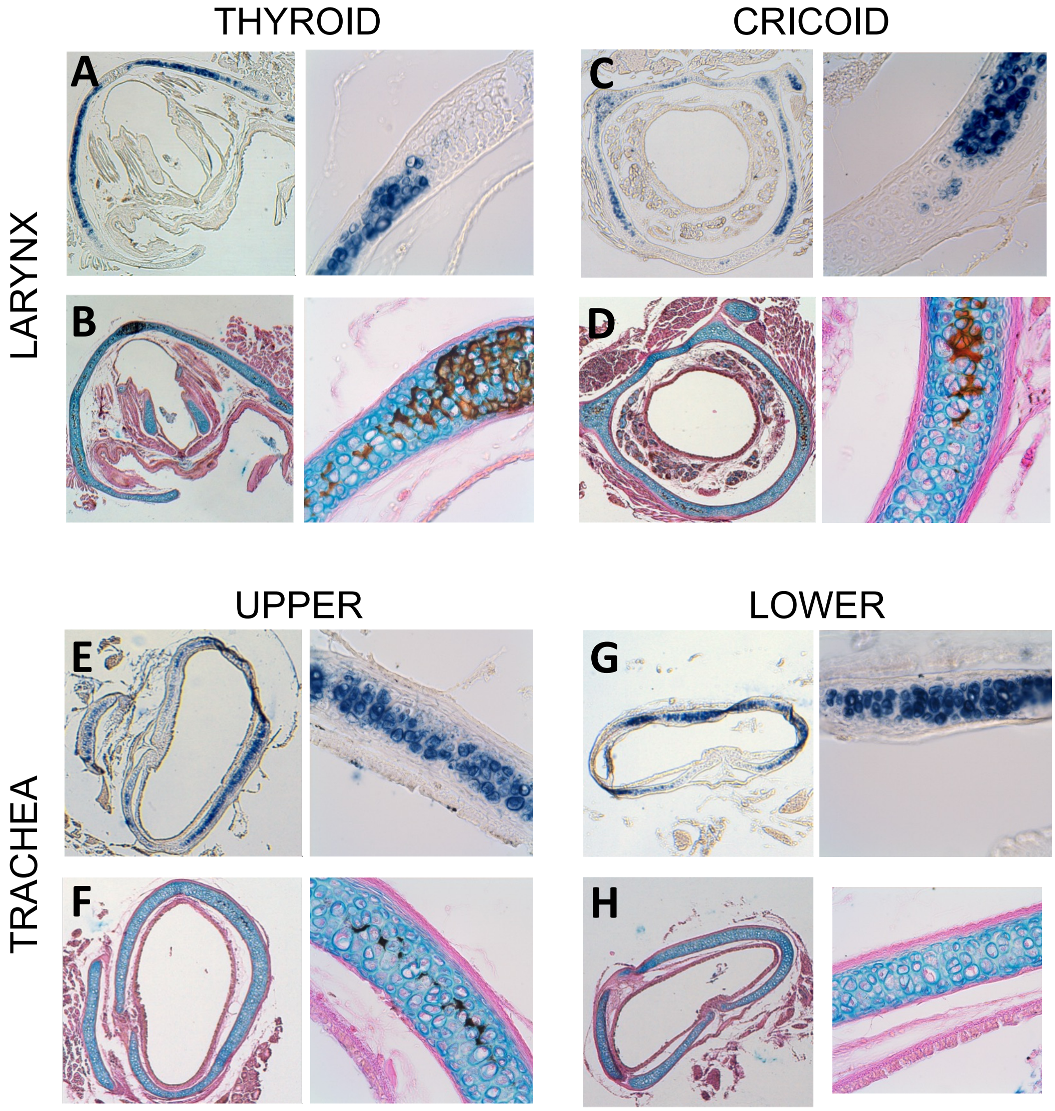
THYROID
CRICOID
LARYNX
A
B
C
D
UPPER
LOWER
TRACHEA
E
F
G
H

### Baptista et al Supp Figure4.pdf

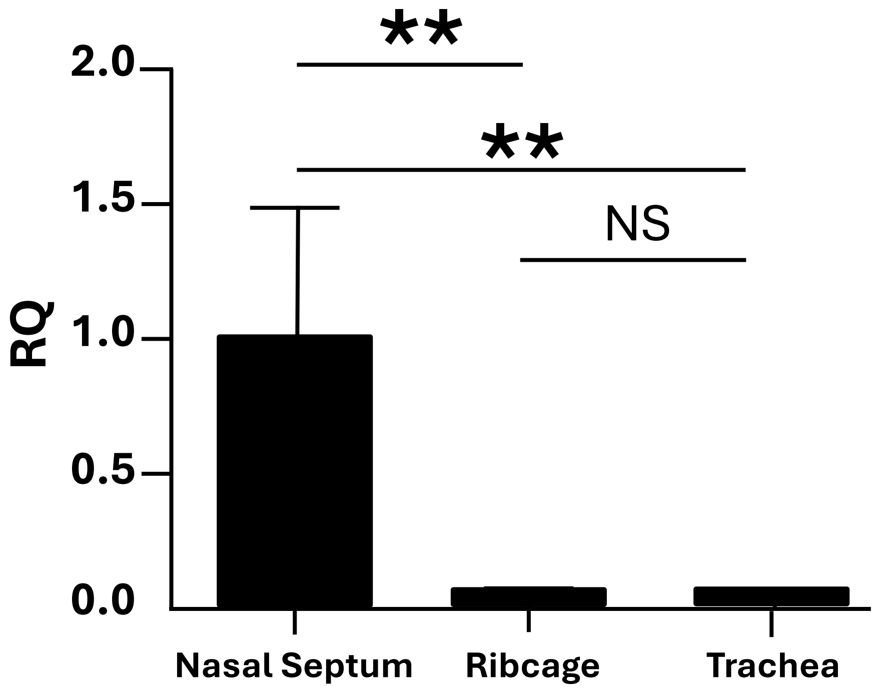
**
**
NS
2.0
1.5
1.0
0.5
0.0
RQ
Nasal Septum
Ribcage
Trachea
